# Molecular Profiles of Matched Primary and Metastatic Tumor Samples Support a Linear Evolutionary Model of Breast Cancer

**DOI:** 10.1101/667303

**Authors:** Runpu Chen, Steve Goodison, Yijun Sun

## Abstract

The interpretation of accumulating genomic data with respect to tumor evolution and cancer progression requires integrated models. We developed a computational approach that enables the construction of disease progression models using static sample data. Application to breast cancer data revealed a linear, branching evolutionary model with two distinct trajectories for malignant progression. Here, we used the progression model as a foundation to investigate the relationships between matched primary and metastasis breast tumor samples. Mapping paired data onto the model confirmed that molecular breast cancer subtypes can shift during progression, and supported directional tumor evolution through luminal subtypes to increasingly malignant states. Cancer progression modeling through the analysis of available static samples represents a promising breakthrough. Further refinement of a roadmap of breast cancer progression will facilitate the development of improved cancer diagnostics, prognostics and targeted therapeutics.

## Introduction

Human cancer is a dynamic disease that develops over an extended period of time through the accumulation of genetic alterations. Once initiated, the advance of the disease to malignancy can be viewed as a Darwinian, multistep evolutionary process at the cellular level [1, 2]. Delineating the dynamic disease process and identifying pivotal molecular events that drive stepwise disease progression would significantly advance our understanding of tumorigenesis and provide a foundation for the development of improved cancer diagnostics, prognostics and targeted therapeutics. Traditionally, system dynamics is approached through time-series studies achieved by repeated sampling of the same cohort of subjects across an entire biological process. However, due to the need for timely surgical intervention upon diagnosis, it is not ethically feasible to collect time-series data to study human cancer. Consequently, while the concept of cancer evolution has been widely accepted [1, 3], the biological process of how cancer progresses to a malignant, life-threatening disease is still not well-understood. The lack of time-series data has been recognized by the field as the central problem in studying cancer progression [4].

With the rapid development of sequencing technology, many thousands of excised tumor tissue specimens are being collected in large-scale cancer studies. This provides us with a unique opportunity to develop a novel analytical strategy to use static data, instead of time-course data, to study disease dynamics. The strategy is based on the rationale that each excised tissue sample provides a snapshot of the disease process, and if the number of samples is sufficiently large, the genetic footprints of individual samples populate progression trajectories, enabling us to recover disease dynamics by using computational approaches. We developed a comprehensive bioinformatics pipeline [5] and applied it to the gene expression data from over 3,100 breast tumor samples available from the TCGA and METABRIC consortiums [6, 7]. Our analysis demonstrated that it is *indeed* possible to use static sample data to study disease dynamics and led to one of the first working models of breast cancer progression that covers the entire disease process. The progression pattern was confirmed by analysis of a series of smaller independent breast cancer datasets and by aligning established clinical and molecular traits with the model. To further validate the model, here we proposed a novel strategy to investigate the progression relationships of matched primary and metastasis tumor samples from breast cancer patients. Our analysis suggested that while breast cancer is a genetically and clinically heterogeneous disease, tumor samples are distributed on a low-dimensional manifold, that disease subtypes are not hardwired and can shift within the same individual, and that the shift is unidirectional along a continuum of disease state towards malignancy. This study shed light on some longstanding issues regarding the origins of molecular subtypes and their possible progression relationships.

## Results

### Constructing a Progression Model of Breast Cancer

As a foundation for the investigation of the relationships between matched tumor pairs, we applied our computational pipeline [5] to the METABRIC data [7] to construct a progression model of breast cancer. The dataset contains the expression profiles of 25,160 genes obtained from 1,989 surgically excised breast tumor samples. Briefly, we first performed supervised learning by using the breast cancer subtypes as class labels and selected 359 disease related genes. Then, we performed a clustering analysis on the expression measures of the selected genes to detect genetically homogeneous groups. By using gap statistic [8] and consensus clustering [9], ten distinct clusters were identified. Finally, we constructed a progression model and represented it as an undirected graph, by using the centroids of the identified clusters as the vertices and connecting them based on the progression trend inferred from the analysis (See Supplementary Data for details). Fig. 1A-B show the distribution of the tumor samples in a three-dimensional space supported by the selected genes and a schematic of the constructed model (referred to as the METABRIC model hereafter), respectively. Our analysis identified a linear, branching model describing two distinct trajectories to malignancy, either directly to the basal subtype with little deviation, or a stepwise, more indolent path through the luminal subtypes to the HER2+ subtype. The two trajectory termini (i.e., HER2+ and basal) represent the two most aggressive breast tumor subtypes [10]. Significant side-branches are also evident for both luminal A/B subtypes. Our results confirmed that molecular subtypes are not hardwired, and genotypes and phenotypes can shift over time [1], and that cancer development follows limited, common progression paths [6].

**Figure 1:**
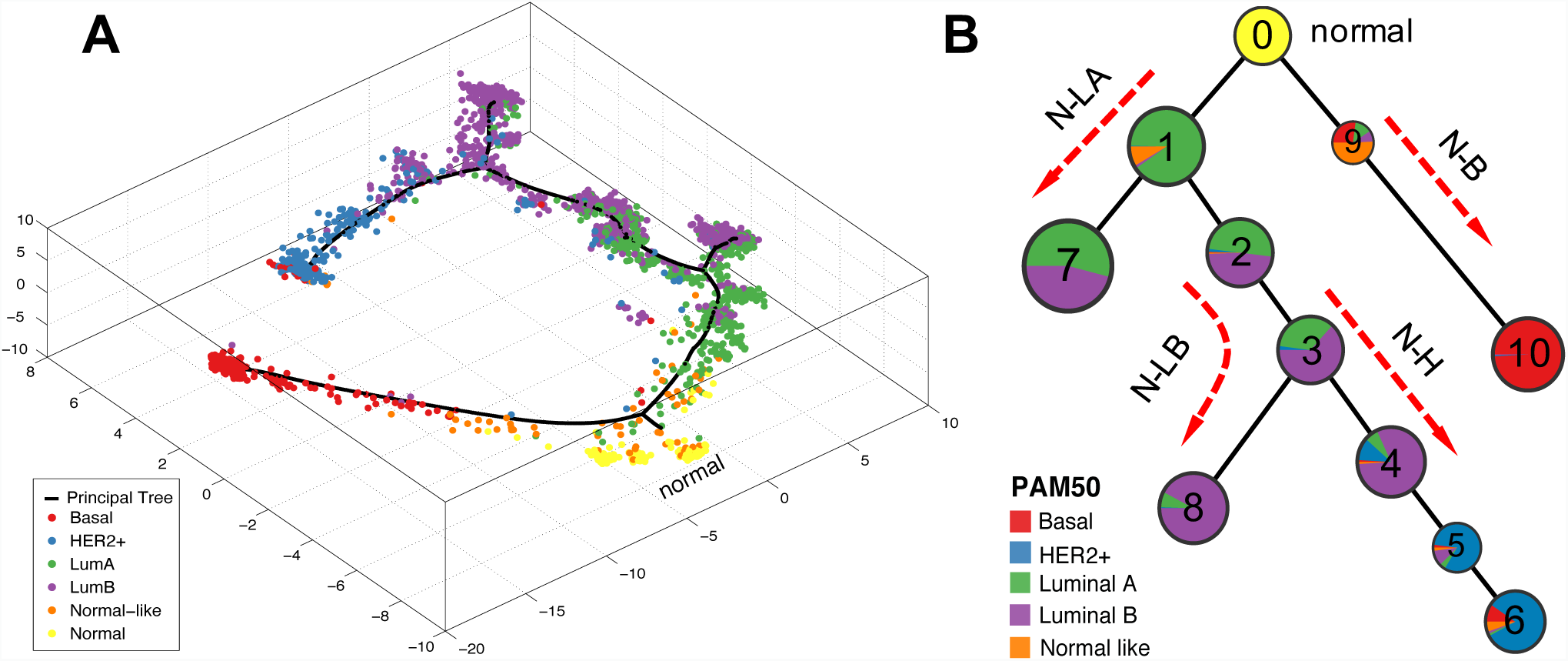
Breast cancer progression modeling analysis performed on METABRIC gene expression data. (A) Data visualization analysis provides a general view of sample distribution. The dataset contains 144 normal breast tissue samples, which we used as the baseline to represent the origin of cancer progression. To help with visualization and to put the result into context by referring to previous classification systems, each sample was color-coded based on its PAM50 subtype label. (B) Constructed progression model of breast cancer. Each node represents an identified cluster, and the pie chart in each node depicts the percentage of the samples in the node belonging to one of the five PAM50 subtypes. The analysis revealed four major progression paths, referred to as N-B (normal to basal), N-H (normal through luminal A/B to HER2+), N-LB (normal through luminal A to luminal B side-branch), and N-LA (normal to luminal A side-branch).

### Mapping of Matched Primary and Metastatic Tumor Samples

To validate the constructed model, we investigated the inter-relationships of 246 matched primary and metastasis (P/M) tumor samples collected from 123 breast cancer patients [11] with respect to molecular subtype and directional progression. As with other RNAseq data, the P/M cohort contains a large amount of missing data and outlier samples, which could complicate the downstream analysis. To address the issue, we performed a series of data pre-processing to remove genes and samples containing excessive missing values, to impute missing data and to identify outlier data (Methods). After data pre-processing, a total of 210 samples including 92 pairs were retained for further analysis. Using the PAM50 classifier [12], we stratified the samples into five intrinsic molecular subtypes, including 3 normal-like, 79 luminal A, 78 luminal B, 33 HER2+ and 17 basal tumors. It has been reported that normal-like samples could be technical artifacts from high contamination of normal tissue [13], so we removed from further analysis the three pairs that contained normal-like classification. Fig. 2A presents a Sankey plot showing the subtype changes of matched pairs.

**Figure 2:**
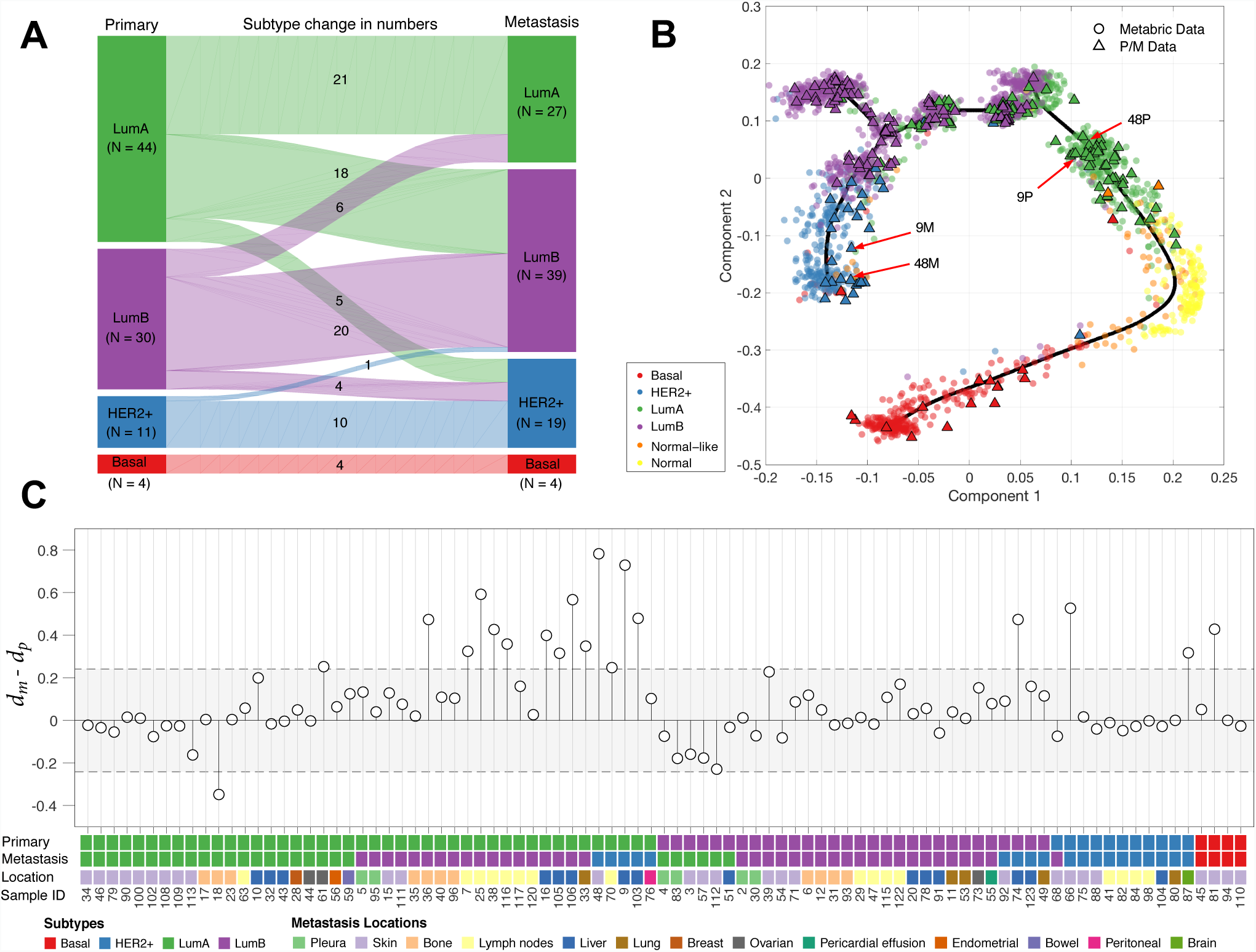
Progression analysis of 89 pairs of matched primary and metastasis (P/M) tumor samples. (A) Sankey diagram showing the subtype changes of matched tumor pairs. (B) Data visualization of the P/M cohort mapped onto the METABRIC model. Two examples (9P/9M, 48P/48M) were shown that underwent evident disease progression from luminal A to the HER2+ subtype. (C) Comparison of progression distances of matched primary and metastasis tumors. A total of 18 and 1 pairs were identified with significant positive and negative disease progression (i.e., samples outside of the shaded region), respectively. *d*_*m*_: progression distance of a metastasis tumor, *d*_*p*_: progression distance of a primary tumor.

We then performed a quantitative analysis of the progression relationships of the primary and metastasis tumors by mapping the data onto the METABRIC model. Since the data sources are not entirely compatible, we first performed batch-effect correction by using ComBat [14] and then applied a novel strategy to map the P/M data onto the 359-dimensional space where the METABRIC model was constructed (Methods). Fig. 2B visualizes the sample distributions of both P/M and METABRIC data, where each sample was color-coded based on its PAM50 label. By using the normal samples as the baseline to represent the origin of cancer progression, we compared the progression distances of each matched pair and reported the results in Fig. 2C. A total of 18 pairs were identified with significant positive disease progression (i.e., the progression distance of a primary tumor is significantly smaller than that of the matched metastasis tumor) and 1 pair with negative disease progression.

From Fig. 2A, we observed that all basal metastatic tumors were derived from basal primary tumors and so were non-basal metastatic tumors. This suggests that basal cancer is a distinct disease entity, supporting the bifurcating structure revealed by the proposed progression model (Fig. 1). We also observed that while most paired samples (62%) were of the same molecular subtype, subtypes did shift over time, primarily from a lesser to a more malignant phenotype (i.e., luminal A to B or to HER2+ subtype), which aligns well with the METABRIC model. There were several examples of luminal B shifting to luminal A (6 pairs) and HER2+ shifting to luminal B (1 pair), implying that cancer evolution can be bi-directional. However, since the PAM50 system provides only an approximate stratification of breast cancer [15], the P/M cohort measured only the expression levels of 87 genes, and RNAseq data contains considerable measurement errors, a qualitative analysis of subtype changes tells only part of the story. Indeed, as shown in Fig. 2B, luminal A and B do not have a clear-cut boundary, as is the case for luminal B and HER2+. This boundary overlap of approximate subtypes explains why in some large-scaled benchmark studies it has been observed that existing molecular subtyping methods only achieve moderate concordance, particularly when classifying luminal A and B tumors [16]. Through a quantitative analysis of the progression distances revealed by mapping the P/M data to the model (Fig. 2B, C), we found that all six luminal B primary tumors that changed to luminal A metastases and one HER2+ primary tumor that changed to a luminal B metastasis had very small disease progression. Conversely, all 18 pairs with significant positive progression were either luminal A to luminal B (9 pairs), luminal A to HER2+ (4), luminal B to HER2+ (3), or basal to basal (2). As examples, Fig. 2B shows the locations of two pairs (9P/9M, 48P/48M) that underwent evident disease progression. Taken together, these analyses support a unidirectional, linear evolution process for breast cancer through luminal subtypes towards malignancy. Interestingly, one tumor pair (18P/18M - both classified as luminal A) had a significant negative disease progression. Although the observed trend was a downstream P-to-M shift, a number of scenarios can coexist. Tumor cells can escape the primary lesion and survive at early stages of primary tumor development. The evolutionary time index may be different for these foci, especially if the metastasis lesion is dormant for a period [17], and local selective pressures at primary and secondary sites can differentially impact the evolution of the related lesions.

## Discussion

Cancer evolution theory dates back to the 1970s [18], and numerous studies have been conducted that significantly expanded our understanding of the concept [1, 2, 4, 19]. Yet, due to the difficulty in obtaining time-series data, beyond conceptual models, there is currently no established cancer progression model derived from tumor tissue data that covers the entire disease process. We have proposed a novel computational strategy that overcomes the existing sampling restrictions. The application of the approach to large-scale breast cancer genomic datasets identified a linear, bifurcating progression model describing two distinct pathways to cancer malignancy. The interpretation of the model is that the basal subtype is distinct from the luminal subtypes, and that the luminal subtypes can shift during disease progression and may be considered as different stages of the same disease. The mapping onto the progression model of the paired primary/metastasis samples further supports the overall model and the concept that the cancer evolutionary process is unidirectional through luminal subtypes towards increasingly malignant states.

Since the proposal of the cancer molecular subtypes [10], fundamental issues regarding whether subtypes are biologically independent entities or have progressive relationships have been under debate [15, 20]. One conceptual model proposes a distinct-path scenario where each subtype follows a path of initiation and progression independently. The alternative is a linear evolution model, where tumors gradually evolve from normal cells to malignant states through the accumulation of genetic alterations [15]. While both models embrace the notion of cancer evolution, the first implies that subtypes are different diseases, while the second suggests that subtypes are different stages of the same disease. Clarifying this issue could have a *profound* impact on current cancer research because clinical management and research strategies in the two scenarios may be very different. Our analysis supports the second model as a representation of disease progression, but also suggests that basal and luminal/HER2+ subtypes are differentially derived from a normal-cell origin.

The development of cancer progression models can inform a range of research directions. For example, current prognostic tests are of value only in a restricted set of patients, but if we can visualize the entire, ordered progressive process, the identification of specific molecular characteristics associated with a broader spectrum of cancer phenotypes becomes feasible. Assisted by genomic testing, we can envisage the placement of samples from individual cases onto a progression path to guide clinical management and evaluate individualized treatment success. The derivation of annotated progression maps can also guide the design of animal studies to focus on pivotal points of cancer development, which may yield the best return with limited resources. Future studies, using higher-resolution genomic methods (e.g., single-cell sequencing and tissue microdissection) and guided by a working model, can provide data for the refinement of a roadmap of breast cancer progression. Although in this study we focus mainly on breast cancer, the developed methods for model construction and validation can also be used to study other cancers and other progressive human diseases, where the lack of longitudinal data is an *unavoidable* problem.

## Materials and Methods

The P/M cohort was downloaded from Gene Expression Omnibus (accession number: GSE92977), which contains the expression levels of 105 breast cancer-related genes and 5 house-keeping genes. The data was log-transformed (base 2) and normalized using the house-keeping genes. To overcome the issue of missing data, we first filtered out 6 genes and 13 samples containing *>* 20% missing values, and then performed missing data imputation on the remaining samples. Specifically, for each sample, we found its 10 nearest neighbors (NNs) and replaced the missing data in each gene in the sample by the average of the observed values of the gene in the identified NNs. The P/M dataset also contains a considerable number of outlier samples, which would complicate downstream analysis. Intuitively, an outlier sample should be dissimilar to its NNs. We exploited this intuition by calculating the average distance between each sample and its 10-NNs and removing 23 samples with top 10% average distances. By using the PAM50 classifier, we stratified the samples into the five intrinsic molecular subtypes (luminal A, luminal B, HER2+, basal and normal-like). With the subtype information, we then performed a one-way ANOVA analysis on individual genes and removed 12 genes that contain little information in discriminating the five molecular subtypes (*p*-value *>* 0.1). After data pre-processing, 87 genes and 210 samples including 92 pairs were retained for the further analysis.

We investigated the progression relationships between the matched tumor pairs by mapping the P/M data onto the METABRIC model. Since the data sources are not entirely compatible, we first mapped the 87 genes in the P/M cohort back to the 25,160 genes in the METABRIC data and identified 85 genes present in both datasets, and then used ComBat [14] to remove the platform-induced data deviations. The METABRIC model was constructed using 359 genes [5] while the P/M cohort contains only the expression data of 85 genes. To address the issue, we designed a novel computational strategy that projects the P/M data onto a high-dimensional space spanned by the 359 genes used to build the METABRIC model. Specifically, we assumed that the projection relationships between the 85 and 359 genes were shared by both cohorts, learned a projection function by using the METABRIC data, and applied the function to the P/M data. See Supplementary Data for a detailed mathematical derivation. After we obtained the high-dimensional prediction of the P/M data, we projected each sample onto the progression paths identified in the METABRIC model. Here, the projection of a sample is defined as a point on a progression path that is the closest to the sample. By using the mean of the normal samples to represent the origin of cancer progression, we compared the progression distances of metastasis tumors with those of their matched primary tumors. Since the P/M data contained considerable measurement errors, we derived a cut-off to identify tumor pairs that underwent significant disease progression (Supplementary Data).

## Supplementary Data

### Bioinformatics Pipeline for Cancer Progression Modeling Analysis

We reconstructed a progression model of breast cancer by applying a bioinformatics pipeline [1] to the data from the METABRIC project [2]. The progression modeling analysis consists of four major steps. First, by using the PAM50 subtypes of individual samples as class labels, we performed supervised learning based feature selection to identify disease related genes. Then, by using the selected genes, we conducted a clustering analysis to identify tumor groups with homogenous genetic profiles. For the purpose of this study, the *K*-means method was employed. The optimal number of clusters was estimated by using gap statistic [3]. It is well known that *K*-means may return a local optimal solution. To identify robust and stable clusters, the technique of resampling-based consensus clustering [4] was employed, where *K*-means clustering was repeated 1,000 times and in each time 80% samples were drawn randomly without replacement from the entire dataset. Then, by using the DDRTree method [5], we constructed a principal tree to mathematically describe the disease dynamics. Finally, by using the constructed principal tree as a backbone, we combined the principal tree and the detected clusters to construct a cancer progression model and extracted disease progression paths. To visualize the sample distribution, we also developed a graph model-based method for dimensionality reduction for structured data [5].

### Progression Analysis of Matched Primary and Metastasis Tumor Samples

The P/M cohort was downloaded from Gene Expression Omnibus (accession number: GSE92977), which contains the expression levels of 105 breast cancer-related genes and 5 house-keeping genes. The data was log-transformed (base 2) and normalized using the house-keeping genes.

#### Data Filtering, Missing Value Imputation and Outlier Removal

As with other RNAseq data, the P/M dataset contains a large amount of missing data. We first filtered out 6 genes and 13 samples containing *>* 20% missing values, and then performed missing data imputation on the remaining samples. Specifically, for each sample, we found its 10 nearest neighbors (NNs) and replaced the missing data in each gene in the sample by the average of the observed values of the corresponding gene in the identified NNs. The P/M dataset also contains a considerable number of outlier samples, which would complicate downstream analysis. Intuitively, an outlier sample should be dissimilar to its NNs. We exploited this intuition by calculating the average distance between each sample and its 10-NNs and removing 23 samples with top 10% average distances.

#### Subtype Classification and Gene Selection

We classified the tumor samples into the five intrinsic molecular subtypes (luminal A, luminal B, HER2+, basal and normal-like), using the PAM50 classifier implemented in the genefu package [6]. A Sankey diagram was generated to visualize the shifts of subtypes between the primary and metastasis tumors. Using the subtype information, we performed a one-way ANOVA analysis on individual genes and removed 12 genes that contain little information in discriminating the five molecular subtypes (*p*-value *>* 0.1). After data pre-processing, 87 genes and 210 samples including 92 pairs were retained for the further analysis.

#### Mapping of Matched Primary and Metastatic Tumor Samples

We performed a quantitative analysis of the progression relationships of the matched primary and metastasis tumors. Given that there are not enough samples in the P/M cohort to build an independent model (a prerequisite of progression modeling analysis is that there are a large number of tumor samples to populate progression paths), we mapped the P/M data onto the METABRIC model. Due to the use of different platforms in generating gene expression data (the METABRIC study used a microarray technique whereas the P/M data was generated through RNAseq), the data sources are not entirely compatible. To remove the platform-induced data deviations, we first mapped the 87 genes in the P/M cohort back to the 25,160 genes in the METABRIC data and identified 85 genes present in both datasets, and then used ComBat [7] to remove batch effect. Note that the METABRIC model was constructed using 359 genes while the P/M cohort contains only the expression data of 85 genes. To address the issue, we designed a novel strategy that projects the P/M data onto a high-dimensional space spanned by the 359 genes used to build the METABRIC model. Specifically, we assumed that the projection relationships between the 85 and 359 genes were shared by both cohorts, learned a projection function by using the METABRIC data, and applied the function to the P/M data.

Let **X** = [**x**_1_, …, **x**_*N*_] and **X**′ = [**x**′_1_, …, **x**′_*N*_] be the METABRIC data in the 85- and 359-dimensional spaces, respectively, and **Y** = [**y**_1_, …, **y**_*M*_] and **Y**′ = [**y**′_1_, …, **y**′_*M*_] be the P/M data in the 85- and 359-dimensional spaces, respectively. We formulated the projection relationship as:

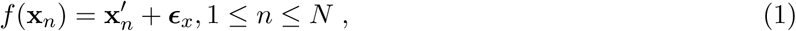

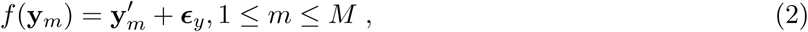

where *f* (·) is a mapping function that projects low-dimensional data to a high-dimensional space, and **ϵ**_*x*_ and **ϵ**_*y*_ are residual terms. In principle, *f* (·) can be a non-linear function representing a complex relationship between two gene sets. However, in this study, we found that a linear form sufficed for our purposes. By using a linear projection, Eqs. (1–2) can be respectively re-written as

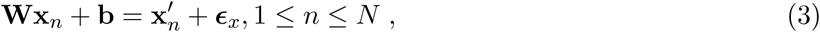

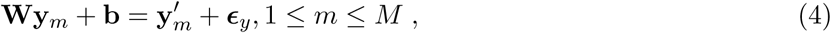

where **W** is a projection matrix and **b** is a bias term, which can be estimated by solving the following least-squares optimization problem:

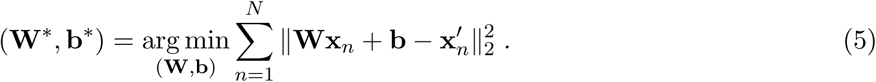

Once we obtained (**W**\*, **b**\*), **Y**′ can be estimated as

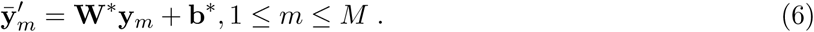

After we obtained the high-dimensional prediction of the P/M data, we projected each sample onto the progression paths identified in the METABRIC model. Here, the projection of a sample is defined as a point on a progression path that is the closest to the sample. By using the mean of the normal samples to represent the origin of cancer progression, we compared the progression distances of metastasis tumors with those of their matched primary tumors. We noticed that many pairs had small differences in their progression distances, likely due to random variations considering that the P/M cohort measured only the expression levels of 87 genes and contained considerable measurement errors. By assuming that the differences in progression distances of tumor pairs that experienced little disease progression follow a zero-mean normal distribution, we estimated the trimmed standard deviation of the differences using the values between 10% and 90% quantiles and identified tumor pairs with a distance difference outside of the ±2*σ* band as those undergoing significant disease progression.

